# Nrf2 Activator PB125^®^ as a Potential Therapeutic Agent Against COVID-19

**DOI:** 10.1101/2020.05.16.099788

**Authors:** Joe M. McCord, Brooks M. Hybertson, Adela Cota-Gomez, Bifeng Gao

## Abstract

Nrf2 is a transcription factor that regulates cellular redox balance and the expression of a wide array of genes involved in immunity and inflammation, including antiviral actions. Nrf2 activity declines with age, making the elderly more susceptible to oxidative stress-mediated diseases, which include type 2 diabetes, chronic inflammation, and viral infections. Published evidence suggests that Nrf2 activity may regulate important mechanisms affecting viral susceptibility and replication. We examined gene expression levels by GeneChip microarray and by RNA-seq assays. We found that the potent Nrf2 activating composition PB125® downregulates ACE2 and TMPRSS2 mRNA expression in human liver-derived HepG2 cells. ACE2 is a surface receptor and TMPRSS2 activates the spike protein for SARS-Cov-2 entry into host cells. Furthermore, in endotoxin-stimulated primary human pulmonary artery endothelial cells we report the marked downregulation by PB125 of 36 genes encoding cytokines. These include IL1-beta, IL6, TNF-α, the cell adhesion molecules ICAM1, VCAM1, and E-selectin, and a group of IFN-γ-induced genes. Many of these cytokines have been specifically identified in the “cytokine storm” observed in fatal cases of COVID-19, suggesting that Nrf2 activation may significantly decrease the intensity of the storm.

## 1. Introduction

Nrf2, is a transcription factor encoded in humans by the NFE2L2 gene. It has been called by many the “master regulator of cellular redox homeostasis” [1], as well as the “guardian of healthspan” and “gatekeeper of species longevity” [2]. Nrf2 regulates most of the genes that defends us against oxidative stress, including superoxide dismutases, catalase, numerous peroxidases, and glutathione metabolism, as well as hundreds of genes involved in scores of important metabolic pathways [3]. The individual vulnerabilities of various structural and catalytic gene products to damage or inactivation by reactive oxygen species (ROS) may lead to degenerative diseases and metabolic dysfunctions. Importantly, Nrf2 declines with age [4,5] and contributes greatly to the “frailty” associated with aging [6-8]. Because Nrf2 transcriptionally upregulates genes that combat oxidative stress, its loss allows oxidative stress to go unmitigated and drive the aging phenotype [1,8,9]. Oxidative stress is therefore a common theme among the key features associated with the aging process, collectively referred to as the “hallmarks of aging”, as it disrupts proteostasis [10], alters genomic stability [11], alters susceptibility to viral and microbial infections [12], and leads to cell death. It is this age-related frailty [8] that often defines the most vulnerable population in situations such as the one we currently face with the coronavirus pandemic [13].

A number of published studies have implicated Nrf2 as a regulator of susceptibility to respiratory viral infections. A recent review by Lee [14] points out that virus-induced modulation of the host antioxidative response has turned out to be a crucial determinant for the progression of several viral diseases. A virus needs to keep oxidative stress at a level optimal for viral reproduction, which is higher than normal, to support the viral metabolism but should not be so high as to kill off the host cell. Viruses have evolved mechanisms for manipulating the Nrf2 pathway in both directions, depending on the needs of the virus, but importantly taking control away from the host cell. Among the types of virus studied are influenza virus, respiratory syncytial virus (RSV), and human metapneumovirus (hMPV) [12,15-18]. The phenomenon is also seen in non-respiratory viruses including Dengue virus (DENV) [19], rotavirus [20], herpes simplex virus [21], Zika virus [22], and HIV [23], suggesting that regulation of oxidative stress may be a need common to most, if not all viruses, and that Nrf2 activators may offer multiple ways to regain control of important pathways to increase resistance and slow viral replication. We recently described a phytochemical composition, PB125, that potently activates Nrf2 by controlling multiple steps involved in the process, especially via the Akt1/PI3K/GSK3β/Fyn pathway [24].

The purpose of this study was to evaluate the effects of Nrf2 activation via PB125 on human liver-derived HepG2 cells as well as on primary human pulmonary artery endothelial cells (HPAEC) in culture. The endothelial cell has been recently implicated as a major player in the tissue destruction caused by COVID-19. Varga et al. performed post-mortem analysis of COVID-19 patients finding involvement of endothelial cells across vascular beds of multiple organs, including electron microscopic evidence of virus particles in renal endothelial cells [25]. Ackermann et al. also documented virus particles inside the cells. Moreover, they found the lungs of Covid-19 patients had a distorted vascularity with structurally deformed capillaries that showed sudden changes in diameter and the presence of intussusceptive pillars that might be explained by drastically dysregulated angiogenesis [26]. These findings broaden the focus of COVID-19 from a disease of pulmonary epithelium to one of multi-organ vascular endothelium. Thus, we measured the expression of genes known to be important for antiviral activity in general, as well as genes with specific relevance to COVID-19 such as ACE2 and TMPRSS2 which determine whether cell types are susceptible to viral entry [27], HDAC5 which helps maintain Nrf2 in an activated state [28], plasminogen (PLG) which is newly recognized as a regulator of cytokine signaling [29], and tissue plasminogen activator inhibitor, PAI-1 (SERPINE1) which has recently been shown to play an important role in the inhibition of host proteases (including TMPRSS2) responsible for influenza A virus maturation and spread [30]. In addition, a group of 36 cytokines expressed by endothelial cells was significantly downregulated suggesting that PB125 might be useful in attenuating the over-exuberant production of cytokines known as cytokine release syndrome or a “cytokine storm” that characterizes a small group of hyperinflammatory conditions that includes graft versus host disease [31], acute respiratory distress syndrome (ARDS) and COVID-19 [32].

## 2. Materials and Methods

### 2.1. Materials and Reagents

Plant extracts: rosemary extract from *Rosmarinus officinalis* (standardized to 6% carnosol; 15% carnosic acid) was obtained from Flavex (Rehlingen, Germany), ashwagandha extract from *Withania somnifera* (standardized to 2% withaferin A) was obtained from Verdure Sciences (Noblesville, IN, USA), and luteolin (standardized to 98% luteolin, from *Sophora japonica*) was obtained from Jiaherb (Pine Brook, NJ, USA). For making PB125 solutions, the rosemary, ashwagandha, and luteolin powders were mixed at a 15:5:2 ratio by mass, then extracted at 50 mg of mixed powder per mL in ethanol overnight and the supernatant isolated [24]. Cell culture: media and antibiotics were purchased from Thermo Fisher Scientific (Waltham, MA, USA). LPS (lipopolysaccharides from Escherichia coli O55:B5) was from Sigma-Aldrich (St. Louis, MO, USA).

### 2.2. Cell Culture

We utilized the human HepG2 cell line (hepatocellular carcinoma) and primary human pulmonary artery endothelial cells (HPAEC) for genomic assays. HepG2 and HPAEC cells are suitable models in the present work because they each have a Nrf2 pathway that responds in a normal manner to Nrf2 activators [33,34], and do not have reported mutations in Nrf2/KEAP1. The HepG2 cells were cultured and maintained by standard methods, using Opti-MEM medium with 4% fetal bovine serum (FBS) and geneticin/penicillin/streptomycin. HPAEC cells were procured from Lonza (catalog # CC-2530) and cultured in Endothelial Basal Media-2 (Lonza catalog #: CC-3516) supplemented with endothelial growth factors optimized for aortic and pulmonary arterial endothelial cells (Lonza catalog # CC-3162). HPAEC subculturing was limited to six passages in order to prevent senescence and de-differentiation. HPAEC were seeded at a density of 5 × 10^5^ cells per 100 mm tissue culture dishes and incubated at 37°C and 6.5% CO2 to 80-90% confluence. All experiments were performed with HPAEC at 80-90% confluence.

### 2.3. IL-6 Protein Assay

We used the Human IL-6 Quantiglo ELISA (R&D Systems, Minneapolis, MN) according to the manufacturer’s instructions to determine the concentration of IL-6 protein released from HPAEC cultured under various conditions.

### 2.4. Gene Expression Assays

#### 2.4.1. Cell Culture and RNA Isolation

To examine the effects of PB125 on gene expression in HepG2 cells, the cells were subcultured in 24-well plates then treated overnight with 0 (control) or 16 μg/mL PB125 (as a 50 mg/mL extract in 100% ethanol). To examine the effects of PB125 on genes that are induced by endotoxin exposure and which may contribute to the cytokine storm (as is observed in COVID-19 illness), we examined a model of pro-inflammatory lipopolysaccharide (LPS) treated human pulmonary arterial endothelial cells, with and without treatment with PB125. HPAEC cells were cultured overnight in 24-well plates under four conditions: control (untreated); PB125-treated (at 5 µg/mL); LPS-treated (at 20 ng/mL); and PB125 + LPS treated. Cells were washed twice with PBS, then extracted with Trizol for total RNA isolation. Further purified with Qiagen RNeasy clean-up columns (QIAGEN Inc., Valencia, CA, USA) as previously described [24].

#### 2.4.2. Microarray Assays

For each sample RNA concentration was determined by absorbance at 260 nm with a NanoDrop spectrophotometer (Thermo Fisher Scientific, Waltham, MA, USA). RNA quality was assessed by Agilent TapeStation 2200 (Agilent, Santa Clara, CA, USA). Gene expressions were determined by the University of Colorado AMC Genomics and Microarray Core facility (Aurora, CO, USA). The GeneChip 3’ IVT PLUS Reagent Kit (Affymetrix/Thermo Fisher Scientific, Waltham, MA, USA) was used to convert 150 ng of total RNA to cDNA according to the manufacturer’s protocol. Each labeled sample was assayed with the Affymetrix PrimeView human gene expression array read with an Affymetrix GeneChip Scanner 3000 (Affymetrix/Thermo Fisher Scientific, Waltham, MA, USA).

The gene transcript and variants are examined using 9–11 perfectly matched (PM) probes. The intensity of expression for all genes on the microarray were evaluated using Affymetrix GeneChip software (Affymetrix/Thermo Fisher Scientific, Waltham, MA, USA) which supported pair-wise comparison between microarray chips.

#### 2.4.3. RNA-seq Library Preparation, Sequencing, and Profiling

Illumina HiSeq libraries (4 assays based on 4 biological replicates in each treatment group) were prepared from HepG2 cell samples using 200–500 ng of total RNA following the manufacturer’s instructions for the TruSeq RNA kit (Illumina, San Diego, CA, USA). With this kit, mRNA is first isolated from total RNA using polyA selection, then the mRNA is fragmented and primed for creation of double-stranded cDNA fragments. Following this, the cDNA fragments are amplified, selected by size, and purified for cluster generation. Subsequently, the mRNA template libraries were sequenced on the Illumina HiSeq 4000 platform (Illumina, San Diego, CA, USA) with single pass 50 bp reads at the University of Colorado Anschutz Medical Center Genomics and Microarray Core Facility (Aurora, CO, USA). Samples were sequenced at a depth to provide approximately 40M single pass 50 bases reads per sample. The derived sequences were then analyzed with a custom computational pipeline comprising the open-source GSNAP [35] Cufflinks [36] and R for sequence alignment and determination of differential gene expression [37]. Reads generated were mapped by GSNAP [35] to the human genome (GRCH38), expression (FPKM) derived by Cufflinks [36], and differential expression analyzed with ANOVA in R.

### 2.5. Statistical Analysis

The data are presented as mean ± standard error of the mean (SEM). One-way ANOVA with Tukey multiple comparisons testing or Student’s *t* test for unpaired data were performed using Prism software (version 6.0, GraphPad Software, San Diego, CA, USA). Statistical significance was set at *p* value < 0.05.

## 3. Results

### 3.1. IL-6 Protein Release

Using the ELISA assay for IL-6, we determined that pretreatment of HPAEC with PB125 decreased the LPS-induced release of IL-6 protein from the HPAEC cells. In this study, the HPAEC cells were plated as described above, then after 24 h they were treated with 5 ug/mL of PB125 extract or with the corresponding amounts of vehicle control. After an additional 16 h of incubation, the cells were treated by adding 20 ng of LPS (or vehicle control) per mL of medium. Each of the four treatment groups was run in triplicate. After 5 hours of LPS treatment, aliquots of cell culture medium were removed from each well for IL-6 measurement by ELISA. LPS stimulation of the vehicle-pretreated HPAEC greatly increased the release of IL-6 protein in to the culture media, but this LPS-induced IL-6 release was reduced by 61% in the cells pretreated with PB125 (*p* = 0.0067). The results are shown in Figure 1.

**Figure 1.**
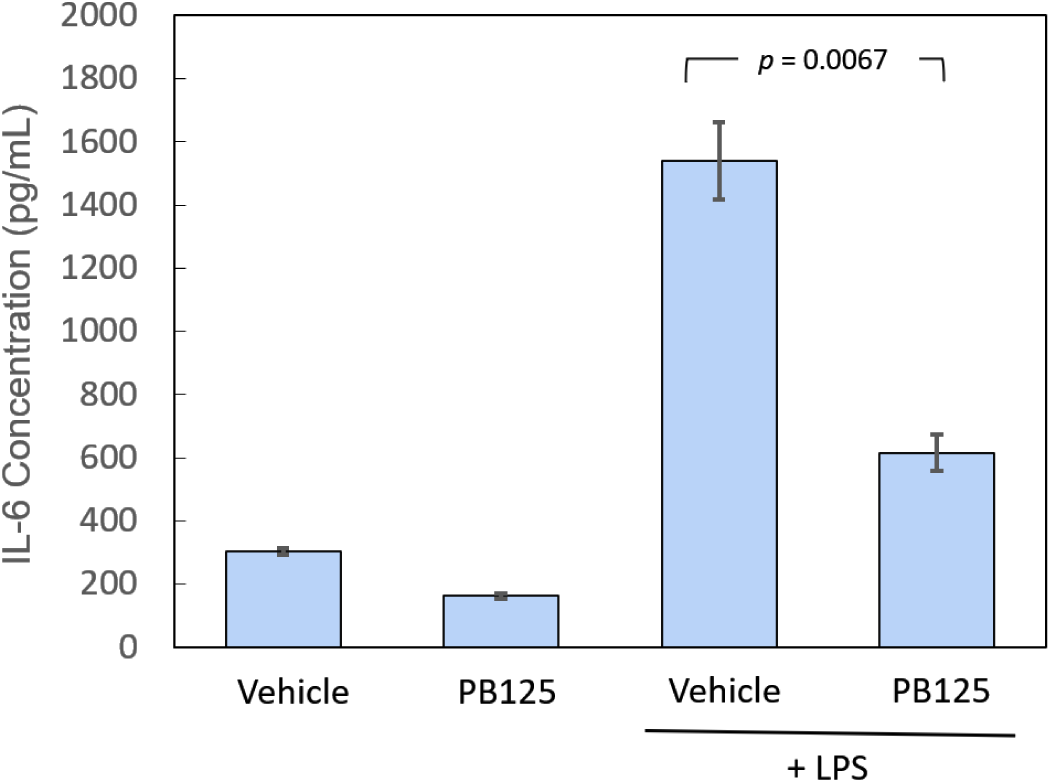
IL-6 protein release is attenuated by Nrf2 activation. HPAEC pretreated with 5 ug/mL PB125, then stimulated 5h with 20 ng/mL LPS had significantly lower levels of IL-6 released into the culture media than vehicle-pretreated HPAEC stimulated with LPS (n = 3 in each group).

### 3.2. Gene Expression

#### 3.2.1. HepG2 Gene Expression by RNA-seq

Because SARS-CoV-2 entry into a human cell depends on ACE2 for binding and on TMPRSS2 for proteolytic activation of the spike protein [27], we examined the effects of PB125 on the expression of these two genes. Because inhibition of the protease activity of TMPRSS2 has been shown to block viral entry [27], we also examined the expression of plasminogen activator inhibitor-1 (PAI-1, encoded by the SERPINE1 gene), a normal plasma component and known potent inhibitor of TMPRSS2 [30]. ACE2 mRNA was down regulated −3.5-fold and TMPRSS2 was down-regulated - 2.8-fold by PB125 in human liver-derived HepG2 cells, as seen in Figure 2. While these impediments may not completely block viral entry, they may significantly impair it, slowing the rate of viral progression. Furthermore, PB125 strongly up regulated SERPINE1/PAI-1 by 17.8-fold. PB125 down-regulated HDAC5 in human liver cells by −2.8-fold, also shown in Figure 2. In humans, HDAC5 appears to be responsible for the deacetylation and attenuation of Nrf2 activity [28]. The cytokine LIF, an important antiviral cellular response to viral infection [38,39], was up regulated 6.6-fold by PB125. Because of recent evidence that plasmin can trigger substantial proinflammatory release of cytokines [29], we examined the effect of PB125 on plasminogen (PLG) mRNA expression, finding it to be down regulated by −1.9-fold. Thus, all six of these gene regulatory effects of PB125 would appear to counter viral attempts to enter the cell and/or to usurp control of oxidative stress response.

**Figure 2.**
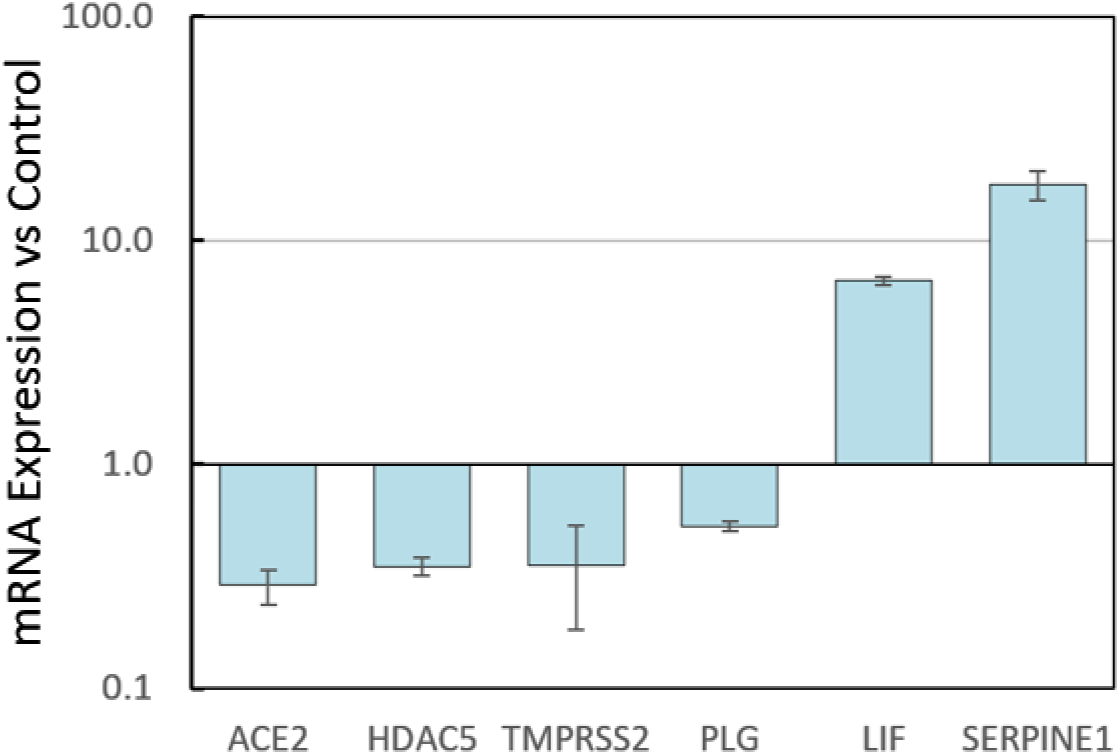
Regulation of pro- and anti-viral genes by PB125. HepG2 cells were cultured overnight in 24-well plates with control vs. 16 µg/mL PB125 and gene expressions were determined using RNA-seq analysis on 4 biological replicates. All six genes differed from control by *p* < 0.04.

#### 3.2.2. HPAEC Gene Expression by Microarray

To examine the effects of PB125 on genes that may contribute to the COVID-19-induced cytokine storm, we examined a model of lipopolysaccharide (LPS) treated HPAEC, with and without treatment with PB125. The results are seen in Figure 3. All 36 genes were significantly upregulated by LPS and normalized to 100% indicated by the red bar (no PB125). Sixteen cytokines, including two colony stimulating factors, are shown in green, with mRNAs downregulated by PB125 as indicated. The average percent inhibition for the group of cytokines was 70%. Two proinflammatory interleukins, IL-1B and IL-6, showed mRNAs inhibited 61% and 44%, respectively. Three proinflammatory cytokine-induced adhesion molecules, intercellular adhesion molecule 1 (ICAM1), vascular cell adhesion molecule 1 (VCAM1), and endothelial cell selectin (SELE) were suppressed an average of 78%. Tumor necrosis factor, TNF, mRNA was reduced by 33%, but a group of five TNF-induced proteins (TNFAIPs) were repressed even more, averaging 70%. Four other genes representing the TNF family were downregulated an average of 65%. Also, a family of five interferon-inducible genes are shown to be downregulated an average of 63%. This across-the-board reduction of genes that contribute to the “cytokine storm” is noteworthy as it is the intensity of this storm that predicts ICU fatalities from COVID-19 [32]. Transforming this storm into a manageable “shower” is therefore a major therapeutic objective in the clinical management of COVID-19 patients.

**Figure 3.**
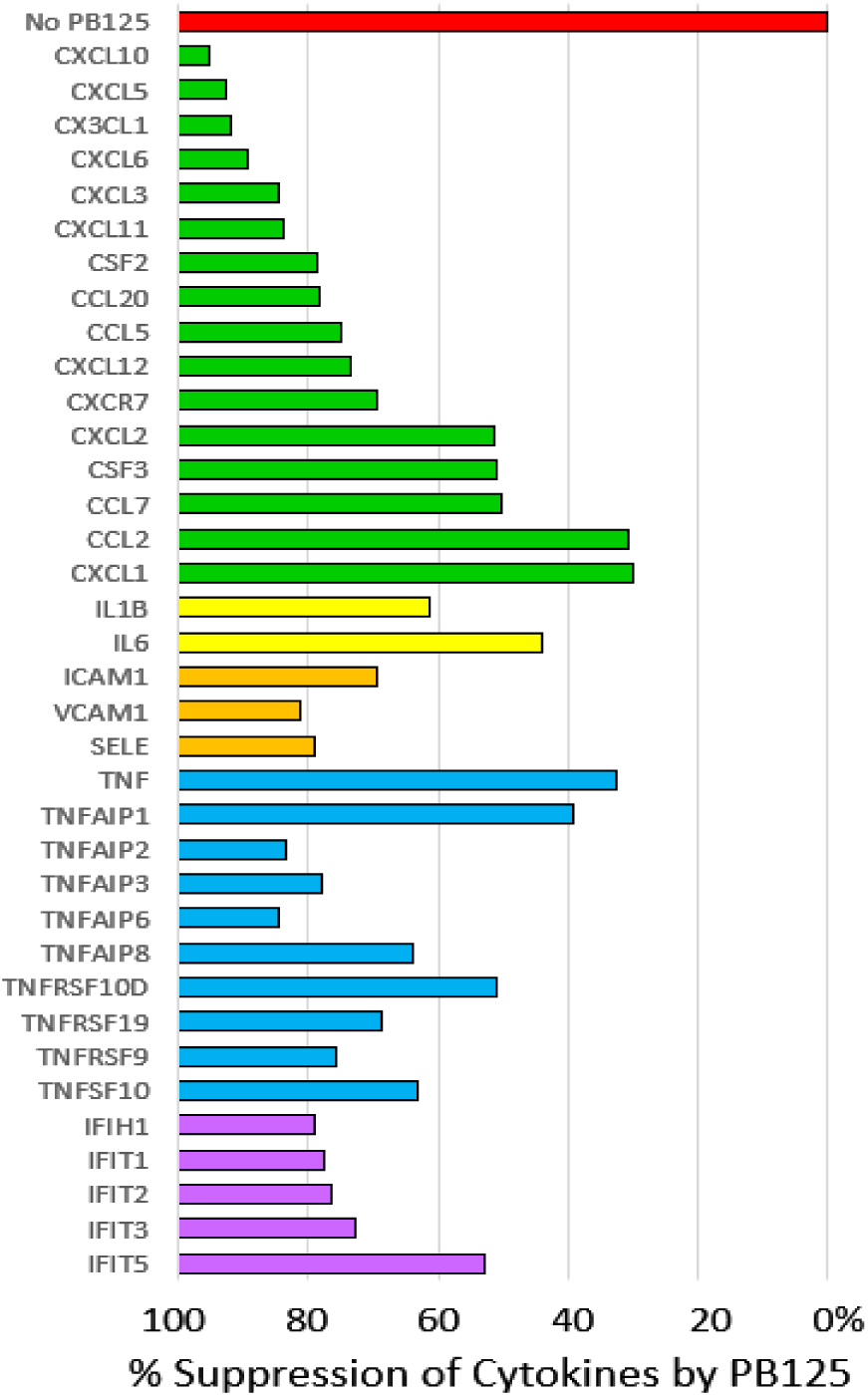
The expression of 36 LPS-induced cytokines in cultured HPAEC was strongly inhibited by PB125 at 5 ug/ml in culture medium. Control expressions were normalized to 100% expression (0% suppression).

## 4. Discussion

The last decade has seen more than 10,000 publications on Nrf2 and its involvement in redox homeostasis, inflammation and immunity, neurodegeneration, aging and diseases associated with aging, ischemia-reperfusion injury, and many other areas, but relatively little has been published regarding its roles in viral infectivity and resistance, despite some rather tantalizing studies. Kesic et al. showed that siRNA knockdown of Nrf2 expression in human nasal epithelial cells effectively decreased both Nrf2 mRNA and Nrf2 protein expression in these cells, which correlated with significantly increased entry of influenza A/Bangkok/1/79 (H3N2 serotype) and replication in the transduced human cells [12]. Importantly, they also demonstrated that enhancing Nrf2 activation via supplementation with sulforaphane (SFN) and epigallocatechin gallate (EGCG) increased antiviral mediators in the absence of viral infection and also abrogated viral entry. Yegeta et al. took a different approach and increased oxidative stress, not just by manipulating Nrf2 expression genetically, but by alternatively exposing mice to an exogenous oxidative stress—cigarette smoke [15]. Cigarette smoke-exposed Nrf2-deficient mice showed higher rates of mortality than did wild-type mice after influenza virus infection, with enhanced peribronchial inflammation, lung permeability damage, and mucus hypersecretion. Cho et al. [40] have similarly studied respiratory syncytial virus (RSV) infection, the single most important virus causing acute respiratory tract infections in children. They found that Nrf2^-/-^ mice infected with RSV showed significantly increased bronchopulmonary inflammation, epithelial injury, and mucus cell metaplasia as well as nasal epithelial injury when compared to similarly infected Nrf2^+/+^ WT mice. The Nrf2^-/-^ mice also showed significantly attenuated viral clearance and IFN-γ, and greater weight loss. Importantly, pretreatment with oral sulforaphane significantly limited lung RSV replication and virus-induced inflammation in Nrf2^+/+^ WT mice. Komaravelli et al. [16-18] have noted that RSV not only causes increased production of ROS *but actively lowers expression of antioxidant enzymes by increasing the rate of proteasomal degradation of Nrf2*. At 6 h post-infection Nrf2-dependent gene transcription was increased, indicating that the cell is in control and responding to the insult of viral infection and the increase in oxidative stress. By 15 h post-infection, however, the concentration of Nrf2 had dropped significantly to about half its pre-infection level reflecting a change-of-control to favor the virus. To accomplish this RSV had increased Nrf2 ubiquitination, triggering its proteasomal degradation, representing one example by which viruses subvert cellular antioxidant defenses. Taken together, these studies demonstrate the ability of Nrf2 to impede viral entry, slow viral replication, reduce inflammation, weight loss, and mortality, but without providing much detailed insight as to which genes and pathways are involved.

The first challenge facing a virus, and particularly a virus that is jumping from one host species to another, is gaining entry to the cell. There are some fairly common ports of entry such as LDLR and ICAM-1 that lead to endosomal entry and which are shared among various viral families [41]. SARS-CoV-2, however, seems to be largely if not totally restricted to a very specific mode of entry. The unique mode of entry may be the greatest vulnerability for the virus, opening the door to some potentially effective therapies. Substantial evidence suggests that a transmembrane protease encoded by the TMPRSS2 gene plays a critical role in the entry for SARS and MERS coronavirus, for 2013 Asian H7N9 influenza virus, and for several H1N1 subtype influenza A virus infections [27,42-45], suggesting that targeting TMPRSS2 could be a novel antiviral strategy to treat coronavirus [42]. Hoffmann and coworkers found that infection by SARS-CoV-2, the virus responsible for COVID-19, may depend almost exclusively on the host cell factors ACE2 and TMPRSS2 [27]. The spike (S) protein of coronaviruses facilitates viral entry into target cells. Entry depends on binding of the S protein to a cellular receptor, ACE2, which facilitates viral attachment to the surface of target cells. In addition, entry requires S protein “priming” by the cellular protease TMPRSS2, which entails S protein cleavage and allows fusion of viral and cellular membranes (Figure 4). This priming can be blocked by clinically proven protease inhibitors of the TMPRSS2, Camostat mesylate [27,46] and Nafamostat mesylate [47]. Nafamostat is remarkably potent with an IC_50_ in the nanomolar range for blocking cellular entry of MERS-CoV in vitro, but has not been clinically tested as an antiviral in humans. The drug appears not to be available at present in an oral formulation, and because of its lack of specificity there is concern over possible side effects [47]. It may prove suitable for treatment of severe COVID-19 cases.

**Figure 4.**
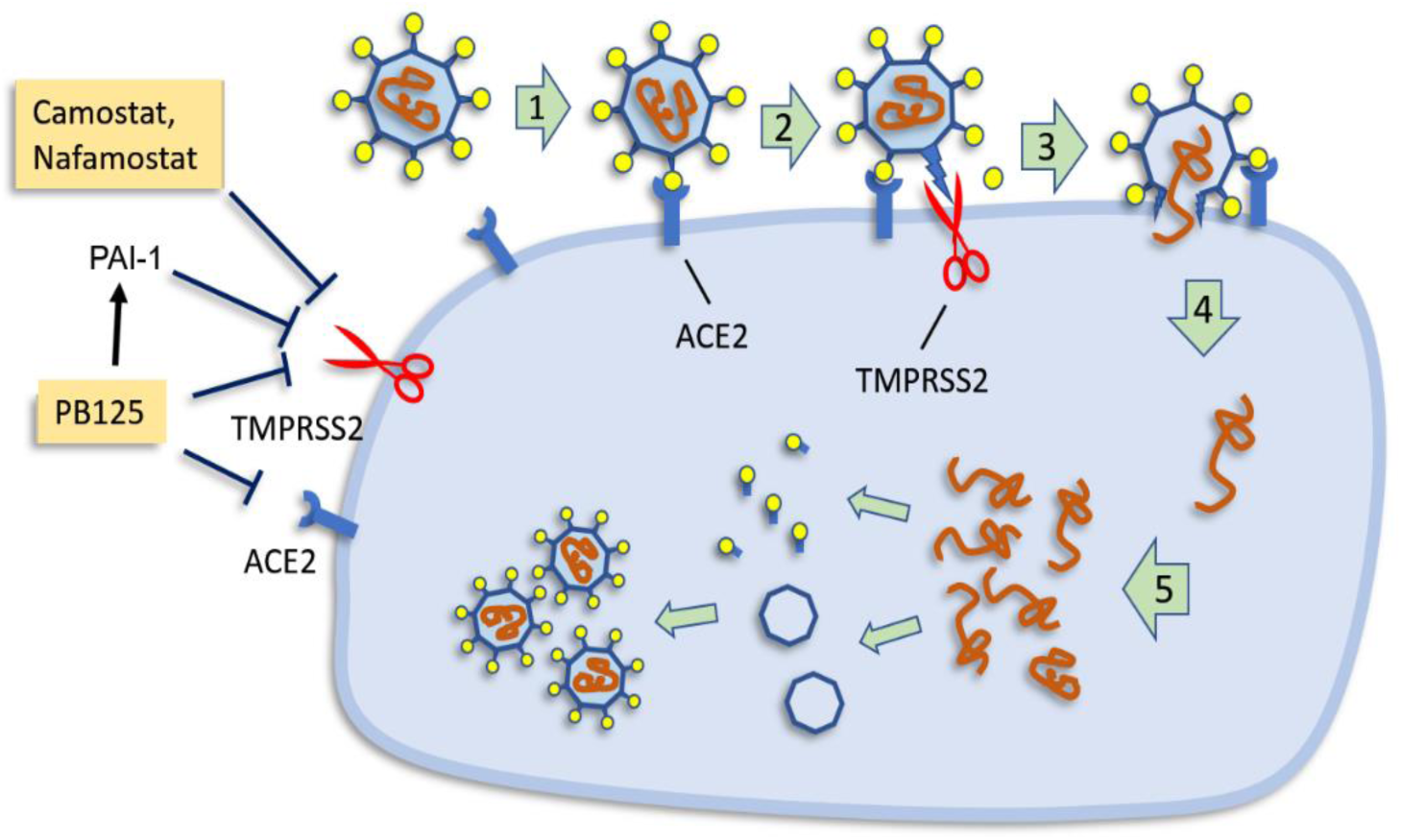
The replication cycle of SARS-CoV-2. Binding of virus to the cell membrane (1) occurs via ACE2 receptors. Spike protein must then be cleaved (indicated as scissors representing serine protease TMPRSS2) to allow entry into the cell (2). The activated spike protein penetrates the cell membrane (3) allowing entry of the viral genome (4), which is replicated, translated, and assembled into mature virus particles (5). On the left, TMPRSS2 inhibition is shown by the antiviral drugs Camostat and Nafamostat, as well as by plasminogen activator inhibitor, PAI-1, encoded by the SERPINE1 gene. PB125 up regulates PAI-2 and downregulates both TMPRSS2 and ACE2 in HepG2 cells.

TMPRSS2 can also be blocked by the human anti-protease Plasminogen Activator Inhibitor-1, or PAI-1 [34], as shown in Figure 4. Iwata-Yoshikawa et al. [43] found that knockout of TMPRSS2 improved both pathology and immunopathology in the bronchi and/or alveoli after infection of the mice by SARS-CoV and completely prevented loss of body weight. This is especially noteworthy in SARS-CoV infection where TMPRSS2 is not the sole mechanism for entry, as is the case with SARS-CoV-2. Thus, our data showing that PB125 down regulated ACE2 by −3.5-fold, down regulated TMPRSS2 by −2.8-fold, and up regulated PAI-1, the potent TMPRSS2 inhibitor, by 17.8-fold (Figure 2) strongly suggest that PB125 treatment might diminish the ability of SARS-CoV-2 to bind to a host cell and to obtain spike protein activation as a result of less ACE2 and TMPRSS2 on the cell surface, and as a result of a 17.8-fold increase in plasma PAI-1, which would inhibit the remaining TMPRSS2. Dittmann et al. [30] have found that influenza A virus (IAV) infection provokes a host response that is both necessary and sufficient for viral inhibition—increased production of PAI-1. They found that for IAV, proteolytic cleavage of the viral coat protein hemagglutinin by host proteases (such as plasmin or TMPRSS2) was a requirement for maturation and infectivity of progeny particles. Addition of recombinant PAI to the apical side of HAEC significantly reduced IAV growth compared to carrier control, with about 90% inhibition of infectivity at 48 h post infection. In contrast, addition of α-PAI-1 antibody dramatically enhanced IAV growth. Thus, many viruses rely on both host endo- and exo-proteases for various entry and maturation functions. Moreover, they found that TMPRSS2, necessary for SARS-Cov-2 infection, is among the trypsin-like proteases effectively inhibited by PAI-1 [30].

A recent publication by Kumar et al. [48] has used computer modeling to predict that withaferin A and withanone, present in extracts of the Ayurvedic plant Ashwagandha (*Withania somnifera*), stably interact at the catalytic site of TMPRSS2, mimicking the pharmaceutical inhibitor Camostat mesylate [27,46,48]. In addition, Kumar et al. found that withanone down regulated the TMPRSS2 gene similarly to what we report here with PB125 (Figure 2). Ashwagandha is one of the three components of PB125, so there is a probability that PB125 may both inhibit the enzymatic activity of TMPRSS2 as well as down regulating the expression of its mRNA.

Quite apart from direct involvement with viral entry mechanisms, PB125 down-regulates HDAC5 in human liver-derived HepG2 cells by −2.8-fold, also shown in Figure 2. Acetylation of Nrf2 increases binding of Nrf2 to its cognate response element in a target gene promoter, and increases Nrf2-dependent transcription of target genes [49]. In humans, HDAC5 appears to be the isozyme responsible for the deacetylation and attenuation of Nrf2 activity [28], and is likely the gene up-regulated by RSV infection described above by Komaravelli et al. [18]. Thus PB125, by inhibiting the deacetylation and subsequent degradation of Nrf2, maintains more active acetylated Nrf2 in the nucleus for a longer time, counteracting one of the mechanisms enumerated above by which viruses attempt to commandeer control of the cell’s redox status and, indirectly, amplifying all Nrf2-dependent actions.

The upregulation of the cytokine LIF, an important antiviral cellular response to viral infection, is shown to be strongly up-regulated by PB125 in Figure 2, again countering potential attempts by a virus to down-regulate it. LIF gene expression was downregulated by H7N9 infection, and knock-down of LIF increased virus titers for three influenza A strains investigated, indicating an important role of LIF in virus defense [39]. In vivo studies were performed with LIF knock-out mice that were infected with RSV. LIF knock-out mice yielded higher virus titers compared to control mice, and LIF signaling was shown to be critical for the protection of the lung from injury during [38].

While we have discussed the role of PAI-1 in preventing TMPRSS2 from activating the SARS-CoV-2 spike protein, mention should also be made of what many would consider its “real job”, the blocking of the conversion of plasminogen (encoded by the PLG gene) to plasmin by both urokinase-type plasminogen activator (uPA) and tissue plasminogen activator (tPA). Fibrinolysis was the first recognized function for plasmin, so one might expect that high expression of PAI-1 would cause low plasmin levels which might be a risk factor for venous thrombosis. Genetic plasminogen deficiency, however, is not strongly associated with risk of thrombosis [50]. In a study of 23 subjects with homozygous mutations in the PLG gene and little or no detectable plasmin, 96% had clinical inflammation of the conjunctivae (ligneous conjunctivitis) but 0% had experienced venous thrombosis [51]. New roles, however, have been recognized for plasmin regarding cytokine release [29,31,52-54]. A “cytokine storm syndrome” is a form of systemic inflammatory response that can be triggered by a variety of factors such as severe infections. It occurs when large numbers of leukocytes are activated and release inflammatory cytokines, which in turn activate yet more leukocytes. Sato et al. found that pharmacological inhibition of plasmin significantly prevented mortality in a mouse model of acute graft-versus-host disease, proposing that plasmin inhibition could offer a novel therapeutic strategy to control the deadly cytokine storm that results from graft-versus-host disease, preventing tissue destruction [31]. Macrophage activation syndrome (MAS) is a life-threatening disorder characterized by a cytokine storm and multiorgan dysfunction due to excessive immune activation. In a mouse model of MAS, Shimazu et al. saw a similar prevention in lethality, concluding that plasmin regulates the influx of inflammatory cells and the production of inflammatory cytokines/chemokines [29]. Plasminogen has also been implicated in activation of astrocytes to produce an array of proinflammatory cytokines [54]. In Figure 2 we report that PB125 downregulates plasminogen mRNA by −1.9-fold in liver cells. Thus, in vivo we speculate that the combined downregulation of plasminogen and 17.8-fold increase of PAI-1 (which covalently reacts with and deactivates both tissue- and urokinase-type plasminogen activators) may significantly attenuate the plasmin-induced cytokine storm phenomenon.

Hyper-inflammation in COVID-19 is associated with such an elevation of proinflammatory cytokines, interleukins, and tumor necrosis factor-α (TNF) and a large number of TNF-induced proteins, and granulocyte colony stimulating factor (GCSF or CSF3). Among 41 hospitalized COVID-19 patients in Wuhan, China, all had elevated IL1B, IP10/CXCL10, and MCP1/CCL2. Sixteen of the patients were subsequently admitted to the ICU and had even higher plasma levels of GCSF/CSF3, IP10/CXCL10, MCP1/CCL2, MIP1A/CCL3, and TNFα, and the intensity of the cytokine storm was a strong prognosticator of mortality [55]. All of these genes encoding these COVID-19-related cytokines appear in Figure 3 as they were significantly up regulated by LPS-treated HPAEC, and significantly down regulated by PB125 treatment. Importantly, a growing number of studies conclude that cytokine storm syndrome is the direct cause of death in most COVID-19 fatalities [55-58].

We speculate that the well-documented age-related loss of Nrf2 expression [4,5] is a potential contributor to the occurrence of a cytokine storm. A longitudinal study of 40 confirmed COVID-19 patients [59] showed that the 13 severe cases, compared to the 27 milder cases, were older (mean age 59.7 vs 43.2), had significantly elevated C-reactive protein (mean 62.9 vs 7.6 mg/l), and showed consistently higher neutrophil counts and lower lymphocyte counts throughout the two week observation period. These observations [55,59] document a clear predilection for severity of COVID-19 infection directly associated with age and intensity of inflammatory response, and presumably inversely associated with Nrf2 expression [8]. The production and self-amplifying nature of an acute inflammatory response demands a prompt subsequent “survival response” from the host tissue to break the self-sustaining attraction of neutrophils to inflamed tissues. We propose that it is a robust oxidative stress-induced activation of Nrf2 in young healthy individuals that follows the gathering storm and rescues host tissues from irretrievable self-inflicted damage. In older individuals, or in the presence of comorbidities that may involve chronic inflammation, the Nrf2-activation response may be insufficient to break the self-perpetuating cycle of events. Figure 5 illustrates this proposed sequence of events. We propose that activating a larger fraction of the limited Nrf2 available in the elderly or otherwise compromised patients might allow them, like their younger counterparts, to shut down cytokine production to stop the escalating cytokine storm, and to begin the recovery and repair phase of the inflammatory episode. This activation boost to suboptimal levels of Nrf2 can be provided by several pharmacological agents, and as well or better by a number of phytochemical activators, as we show here with PB125.

**Figure 5.**
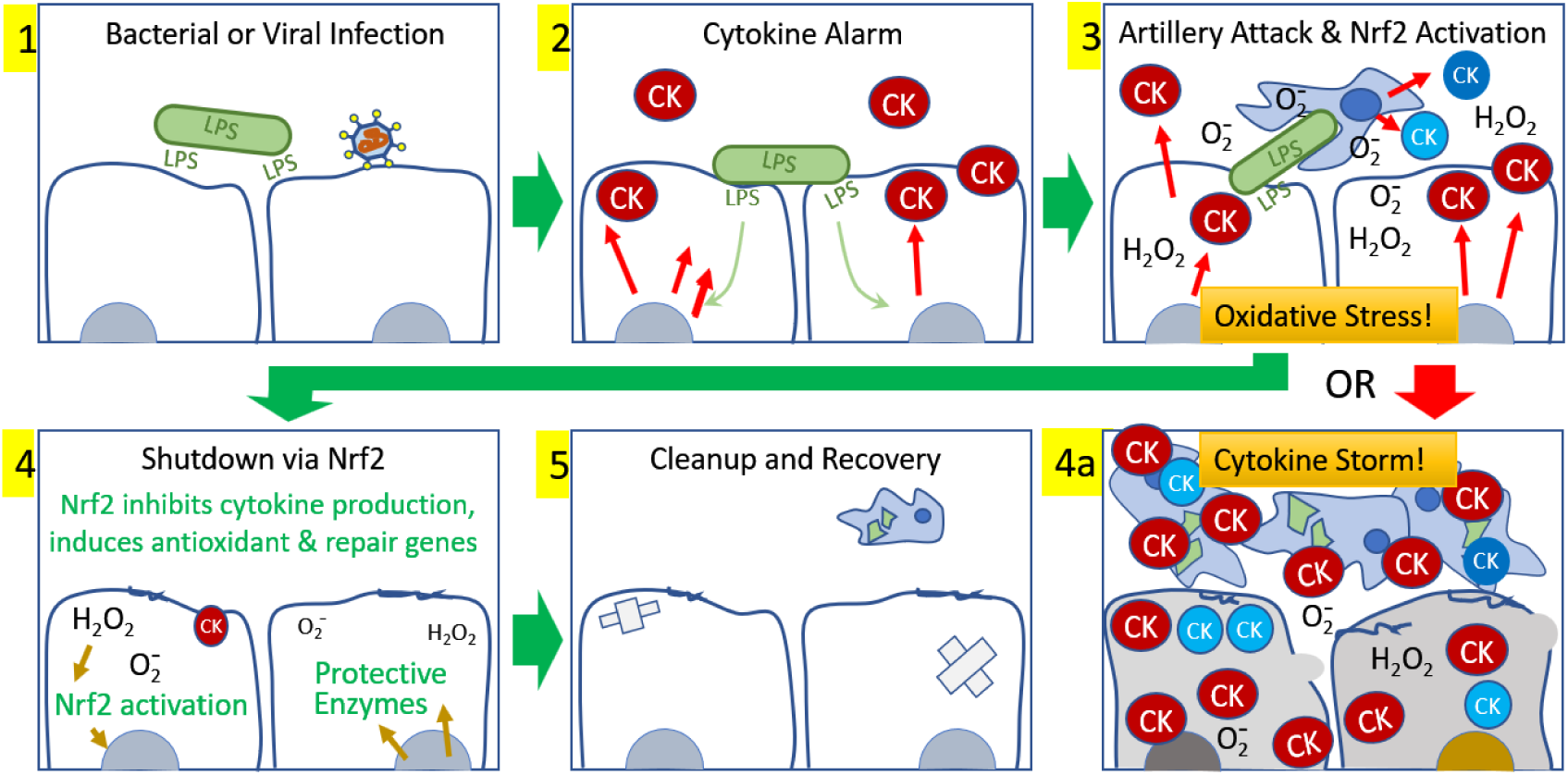
Development and resolution of an acute inflammatory event. 1) Initiation occurs with bacterial or viral infection, which triggers 2) local production of cytokines by endothelial cells to call in inflammatory cells to neutralize the invasion. 3) An attack ensues in which superoxide and secondary oxidants are produced, phagocytosis occurs, and more cytokines are released by the first responders, calling in subsequent waves of activated inflammatory cells. 4) In young healthy cells the oxidative stress generated by the battle activates Nrf2 and within hours the nearby tissues are inhibiting further tissue cytokine production, rescuing host cells from further damage and permitting 5) repair, cleanup, and recovery. Alternatively, in 4a) older cells deficient in Nrf2 may be unable to mount Nrf2 activation sufficient to break the self-sustaining chain reaction, resulting in an uncontrolled cytokine storm that ultimately destroys the host tissue and leads to death. A more robust activation of the limited Nrf2 available in older cells may be provided by pharmacological or phytochemical Nrf2 activators.

It is worth noting that the longitudinal study of COVID-19 patients [59] reported a remarkably higher concentration of serum ferritin in the severe cases (averaging 835.5 μg/l, with a range of 635.4 to 1538.8) versus the milder cases (averaging 367.8 μg/l, with a range of 174.7 to 522.0). Ferritin has long been recognized as a source of iron released under inflammatory conditions by the superoxide radical [60-63]. Under the intense oxidative stress precipitated by a cytokine storm, the release of iron would catalyze lipid peroxidation and greatly amplify host tissue injury. COVID-19 has already been added to the short list of known “hyperferritinemic” diseases, all of which are characterized by high serum ferritin and life threatening hyper-inflammation sustained by a cytokines storm which eventually leads to multi-organ failure [64]. Ferroptosis is a newly described form of regulated cell death that is iron-dependent and causes cell death by mitochondrial dysfunction and toxic lipid peroxidation. Nrf2 has been implicated as a “key deterministic component modulating the onset and outcomes of ferroptotic stress” [65].

We studied three different phenomena that may be involved in vulnerability to the SARS-Cov-2, and these interactive phenomena are not all present in any single cell type. The first objective, to study the effects of PB125 on expression of ACE2 and TMPRSS2, could be studied in presumed point-of-entry cells, including alveolar Type II cells [27] and HPAEC [25,26]. There are both clinical and laboratory suggestions that the virus may invade many organs containing cell types that co-express ACE2 and TMPRSS2 (heart, gut, kidney, eye), including liver, where we showed down regulation of both ACE2 and TMPRSS2. Thus, liver is also a relevant cell in which to study these entry genes. A second objective was to examine whether PB125 could limit systemic plasmin activity by down regulation of PLG and/or upregulation of PAI-1, both plasma proteins not produced by the cells of the lung. In this case, the liver is the source of PLG, and is the appropriate cell type to examine. The source of PAI-1 in plasma isn’t known, but may be liver or muscle. For the third objective of whether PB125 could down regulate the cytokines identified as participating in the cytokine storm phenomenon, we believe the focus on primary HPAEC is appropriate, although a more complete picture would include study of the inflammatory cell types themselves. HPAEC generate the 36 cytokines we examined, sounding the systemic alarm that recruits inflammatory cells to the infected organ. The Type II alveolar cell has very little capacity for producing cytokines [66]. For studying how an intervention can break the inflammatory cycle leading to a cytokine storm, we believe vascular endothelial cells may be the single most important players. A potential limitation of the study is the use of LPS as a surrogate for SARS-CoV-2 to induce the inflammatory response.

We propose that one evolutionary driving force for the Nrf2 pathway may have been to provide a failsafe brake for out-of-control inflammatory events. The increase in oxidative stress at the site of an intense inflammatory locus may have been the intended trigger for a system to activate Nrf2, allowing it to end the assault at a point where the invader has likely been vanquished but from which the host may be able to survive. In fact, the cadre of genes induced by Nrf2 have long been referred to as “survival genes” [3].

## 5. Conclusions

We have shown that a group of 42 genes linked to respiratory virus infectivity and resistance, or to the associated immune response, are responsive to pharmacological Nrf2 activation. It seems possible that the sum total of these multiple antiviral effects may confer a degree of resistance, may attenuate viral replication rate, may alleviate symptoms by limiting microvascular injury, and perhaps allow successful navigation through the “cytokine storm” that is a particular problem with COVID-19. Even though the never-ending evolutionary war-of-wits continues, and the viruses occasionally win a battle, this scenario of the complex and multi-faceted antiviral mechanisms regulated by Nrf2 serves to underscore the importance of this very central transcription factor in keeping us protected and functional.

## Author Contributions

Conceptualization (J.M.M., B.M.H, B.G.); methodology (B.M.H., B.G., A.C.G); investigation (A.C.G., B.G.); data analysis (J.M.M., A.C.G., B.G.); manuscript preparation (J.M.M., B.M.H), manuscript review and editing (J.M.M, B.M.H., A.C.G., B.G.)

## Acknowledgments

The authors thank Ken Jones and Wenhua Ren for their assistance in analyzing the gene array and RNA-seq data. This work utilized the Genomics Shared Resource of the University of Colorado Cancer Center (P30CA046934). The authors dedicate this work to the memory of Kara P. Geraci.

